# Utility of stromal lymphocytes in diagnosis and predicting upgrade of B3 breast lesions from core biopsies

**DOI:** 10.1101/2022.09.02.506444

**Authors:** Tanjina Kader, Shona Hendry, Elena Provenzano, Madawa W Jayawardana, Jia-Min Pang, Kenneth Elder, David J Byrne, Lauren Tjoeka, Helen ML Frazer, Eloise House, Sureshni Jayasinghe, Holly Keane, Anand Murugasu, Neeha Rajan, Islam M Miligy, Andrew R Green, Emad A Rakha, Stephen B Fox, G. Bruce Mann, Ian G Campbell, Kylie L Gorringe

## Abstract

For more than two decades attempts have been made to identify a subset of women diagnosed with lesions with uncertain malignant potential (B3 lesions) who could safely be observed rather than being treated with surgical excision and/or chemoprevention. Various histopathological, clinical and imaging parameters for risk recommendation have been evaluated, with little impact on clinical practice. The primary reason for surgery is to rule out an upgrade lesion to either ductal carcinoma *in situ* (DCIS) or invasive breast cancer (IBC). While on average 30% of these patients are upgraded after diagnostic biopsy, a large number are over treated,making this an important harm of screening.

Here we evaluated stromal lymphocytes from B3 biopsies (n=264) as a predictive biomarker for upgrade. A higher number of stromal lymphocytes were observed in upgraded B3 lesions than non-upgraded (p< 0.01, zero inflated binomial model) for both ductal and papillary lesions (n=174). This observation was validated in an independent cohort (p<0.001, p<0.05, zero binomial model, ductal and papillary lesions, respectively) (n=90). Our data suggested that the presence of ≥5% of lymphocytes in the surrounding specialised stroma of B3 lesions are predictive of B3 lesions being upgraded with a specificity of 93% and 87% in our discovery and validation cohorts, respectively. The area under the curve (AUC) for the discovery cohort using lymphocyte count and age as variables was 0.77 and was validated with an AUC of 0.81 in the validation cohort.

In conclusion, we can identify a subset of the patients at risk of upgrade with high specificity. Assessing the tumour microenvironment including stromal lymphocytes may contribute to reducing unnecessary surgeries in the clinic.

## Introduction

Early detection of malignant breast lesions must be balanced with minimising overtreatment of non-invasive breast conditions in a mammographic screening program [1]. Problematic diagnoses that represent a potential harm of routine mammographic screening are a suite of breast lesions under the general term *lesions of uncertain malignant potential*, also known as B3 lesions [2]. It is estimated that more than 300,000 women are diagnosed with these problematical lesions every year in the United States alone, requiring surgical excision. B3 lesions include atypical ductal hyperplasia (ADH), breast papillary lesions, flat epithelial atypia (FEA), columnar cell change (CCL), radial scar, usual ductal hyperplasia (UDH), sclerosing adenosis and atypical lobular hyperplasia (ALH) [2]. B3 lesions are surgically removed primarily because of the high rate of detecting carcinoma in the excision specimen. Here, we refer to this as “upgrade”, encompassing the scenarios whereby the biopsy missed a nearby carcinoma or there was insufficient indication of invasive breast cancer (IBC) or ductal carcinoma *in situ* (DCIS) in the core biopsy for a definitive diagnosis. Recent meta-analyses suggest an average 29% risk of upgrade for ADH and 36% for papilloma with ADH [3, 4]. Overall, 17% of all B3 lesions are at risk of upgrade [5], leading to overtreatment of many patients.

Misdiagnosis on biopsy further reduces the effectiveness of mammographic screening. Low-risk atypical lesions such as FEA or CCL with atypia can be misdiagnosed as ADH [6]. However, if correctly diagnosed, FEA for example exhibits a very low upgrade rate and life-time risk of developing IBC [7–9]. The similar architectural features of ADH and low grade (LG) DCIS is another diagnostic challenge, and with only the extent of ducts (2 mm size threshold) as a distinguishing feature, ADH bordering on LG DCIS on biopsy confers a high upgrade rate [10, 11]. In addition, recent meta analyses showed that ADH alone can be upgraded to IBC after surgical excision from 9 to 28% of the time [4, 5]. In fact, any B3 lesion with atypia carries a higher risk of missing co-existing malignancy if not excised fully [4, 5]. Thus, such lesions often lead to a recommendation for complete surgical removal [6, 11]. A subset of these tumours can be predicted from mammogram imaging. According to the mammogram imaging category used in this study (i.e. not BIRADS), those with a category 5 (i.e. malignancy) are not problematical lesions in the clinical setting as surgical removal is always indicated. The truly problematical lesions are B3 diagnosed on biopsy with an imaging category 3 (i.e. equivocal) or category 4 (i.e. suspicious for malignancy). Since neither imaging information nor diagnosis on biopsies are sufficient to exclude DCIS including intermediate or high grade (HG) DCIS or any IBC, surgical excision remains the standard of care for all B3 lesions suspected to be ADH in particular [2, 12–14]. There is no robust biomarker to predict upgrade of any B3 lesion, reducing the effectiveness of routine screening.

Tumour infiltrating lymphocytes (TILs) have been evaluated as an independent predictive or prognostic biomarker for IBC [15]. Higher TILs positively correlated with favourable prognosis and overall improved survival in triple negative and HER2 positive IBC [16, 17]. Studies on TILs have shed some light on the immune microenvironment of DCIS and benign breast disease, although the utility of TILs as a recurrence biomarker has not been consistent [18–20]. A very recent study evaluated TILs in DCIS from the COMET trial and found that a higher number of TILs correlated with the upgrade rate of LG DCIS to HG DCIS [21]. Although TILs have clinical utility in breast cancer in a number of different domains with high reproducibility among pathologists [22], they have never been evaluated to predict upgrades of B3 lesions. Here we investigate and report for the first time whether the stromal lymphocytes of B3 lesions could be used to predict upgrade.

## Methods

### Cohorts

For both discovery and validation cohorts, all B3 lesions were divided into 2 histological groups, ductal (ADH/FEA/CCL/radial scar/UDH) and papillary with and without atypia. Based on our previous findings, these two groups, especially ADH and papillary lesions are not only histologically different, but also genetically distinct [23, 24]. All assessed cases in the validation cohort and most cases in the discovery cohort are core biopsy specimens except for fourteen cases from the discovery cohort (Supp File 1). These 14 non-upgraded ADH and papillary lesions were from our previously published studies [23, 24] and were included to have an overall investigation of non-upgraded B3. All cases were treated with surgical excision per standard therapy at the time of diagnosis.

The two discovery cohorts were identified from the Royal Melbourne Hospital (RMH) database (1995-2015) and St Vincent’s Hospital (StVs) database (2000-2016) (n=272) with an addition of 3 upgraded cases from the University of Nottingham, UK. The validation cohort was identified from North West Breast Screen (NWBS) (2004-2013) (n=115). Only cases diagnosed as B3 on biopsies with a subsequent excision (< 1 year) report available in the hospital database were included in this study. Patients with any previous history of breast cancer were excluded (Supp Fig 1, Supp Fig 2). Forty one percent of these cases (160/387) in total had available data for the imaging category from Breast Screen Victoria (BSV) (Supp File 1). The concordance information between pre and post biopsy (i.e. that the needle targeted the exact area that was identified to be biopsied) was also recorded from BSV, confirming all cases were concordant in this study. Therefore, no cases were excluded based on discordance.

### Histopathological criteria for ADH, other ductal B3 lesions and papillary lesions

Diagnostic hematoxylin and eosin (H&E) stained tissue sections were reviewed by a specialist breast anatomical pathologist (N.R., A.M., P.H.) to ensure consistent diagnoses under current criteria. This review process was blinded to the outcome of the subsequent excision (upgraded or not).

#### Criteria for ADH

The criteria for ADH was according to Page *et al*. [25] and Lakhani *et al*. [26] as previously published [23]: “ADH is characterised by a proliferation within TDLU of a monomorphic population of epithelial cells with generally rounded nuclei that are evenly spaced and have well defined cell borders. The cells form “punched out” (cribriform-like) secondary lumens and/or micropapillae. The cells may grow in arcades or rigid bridges of uniform thickness”. The neoplastic cells comprising the proliferation are cytokeratin, CK5/6 negative (surrounding myoepithelial cells show staining for CK5/6) as well as strong and uniform positively stained nuclei for ER as previously published [23]. In distinguishing ADH from LG DCIS, the latter required complete involvement of >2 ducts or partial involvement of ducts >2 mm in extent, in keeping with the criteria described in Lakhani *et al*. [26]. Exclusion criteria of ADH were absence of atypical cells (i.e. UDH), CCL with UDH, and other early neoplasia with atypia such as FEA, radial scar or apocrine hyperplasia. All of these lesions lack the secondary structure of cribriform and/or micropapillary structures, and had only cytological atypia with an architectural structure of CCL or FEA [2, 26] (Supp Figure 3).

Upon re-review, 86/158 initially identified as non-upgraded ADH (discovery cohort) did not meet the criteria of ADH and were other lesions such as FEA, CCL, radial scar and UDH (Supp Figure 2). It should be noted that the initial diagnosis of these cases was carried out between 1995-2016 by multiple different pathologists. With wider use of immunohistochemical markers, the definition and clinical practice for some of these lesions may have changed since then. Even though these non-upgraded ADH cases were not reconfirmed as ADH, they still were included in this study and grouped as “other non-ADH ductal B3 lesions”. However, 20/158 cases that were considered LG DCIS on review were excluded from the upgrade cohort. Only one out of 37 upgraded ADH cases did not meet the criteria and was described as FEA and radial scar. Details of patient selection are in Supp Figure 1 and 2. Molecular characterisation of 20 out of the 53 confirmed non-upgraded ADH cases was previously published in Kader *et al*.[23]. Most of the H&Es from surgical blocks were unavailable and therefore, were not re-reviewed. If upgraded, the type and grade of DCIS or IBC was recorded from their pathology report at the time of diagnosis.

#### Criteria for papillary lesions

The criteria followed for benign papilloma or papilloma with ADH were according to Page *et al*. combined with the recent recommendation of the World Health Organisation (WHO) (papilloma with <3mm extent of ADH) [26, 27]. p63 and CK5/6 immunohistochemistry (IHC) were evaluated when available to determine the differential diagnosis between benign papilloma and papillary carcinoma (p63+ve = benign papilloma, p63-ve = carcinoma) as well as reconfirming the atypical populations (<3mm CK5/6 –ve = papilloma with ADH; >3mm CK5/6-ve = papilloma with DCIS), respectively as previously published [24].

### Assessing stromal lymphocytes on H&E stained sections

The assessment of stromal lymphocytes was based on H&E sections from the core biopsies of any B3 lesions. All were assessed by T.K. Based on the data from the discovery cohort suggesting that B3 biopsies having ≥5% stromal lymphocytes were most likely to be upgraded, the reproducibility of predicting upgraded cases was tested. The reproducibility of a randomly selected 40 cases between observer 1 (T.K.) and observer 2 (S.H.) or observer 3 (J.M.P.) were assessed. Assessors were blinded to the outcome (upgraded/non-upgraded). The cases were called either as <5% or ≥5% stromal lymphocytes and this designation was 100% concordant among the observers.

The method to assess stromal lymphocytes of B3 lesions followed that previously published for assessing TILs of DCIS by Hendry *et al*. [18]. This method was developed by Pruneri *et al*.[19] for DCIS and other pre-malignant lesions, supported by the recent guidelines of the International Immuno-Oncology Biomarker Working group [22]. Very recently the method was also utilised for benign breast disease [20]. Since the B3 lesions are not *tumour*, the term used in this manuscript is stromal lymphocytes instead of TILs.

In brief, the stromal lymphocytes of B3 lesions was reported for the stromal compartment (=% stromal lymphocytes). The stromal component was defined as the area within the “specialised” stroma surrounding the B3 lesions (Figure 1). If the specialised stroma is not clearly visible, the lymphocytes clearly surrounding the B3 lesions should be assessed. The specialised stroma was defined as the area surrounding the duct within two high power microscope fields (∼1 mm). Any minimum, partial; grouped/scattered stromal lymphocytes was taken into account. Stromal lymphocytes of any normal ducts or terminal-ductal lobular unit (TDLUs), if present, were noted while assessing, however, were not taken into account for the total stromal lymphocytes according to the International Immuno-Oncology Biomarker Working Group guidelines [22]. In biopsies with a mix of lesions, such as UDH surrounded by CCL and FEA, the stromal lymphocytes were counted across all lesions and an average score was used. For biopsies with multiple blocks per case, all blocks were assessed and an average lymphocytes count was used. Since the B3 lesions including ADH were diagnosed on core biopsies, and biopsies could be often observed with very limited stromal component, this method was altered slightly for these lesions. Cases were excluded from stromal lymphocyte assessment if there was a lack of stromal components (<1 mm) surrounding the ducts of B3 lesions/ADH/papilloma (Supp Figure 4). This assessment was extremely crucial since many papillary lesions lack the stromal component on biopsies due to the minimal sampling and were excluded. However, most cases met the criteria for ductal B3 lesions and therefore, the total number of assessed cases from the discovery cohort were non-upgraded ADH (n=43), other non-ADH ductal B3 lesions (i.e. FEA, CCL, radial scar etc) (n=55) (Supp Figure 3), papillary lesions (benign and papilloma with ADH) (n=31), upgraded ductal B3 (n=36) and upgraded papillary B3 (n=9) (Supp Figure 1: Patient selection criteria).

**Figure 1.**
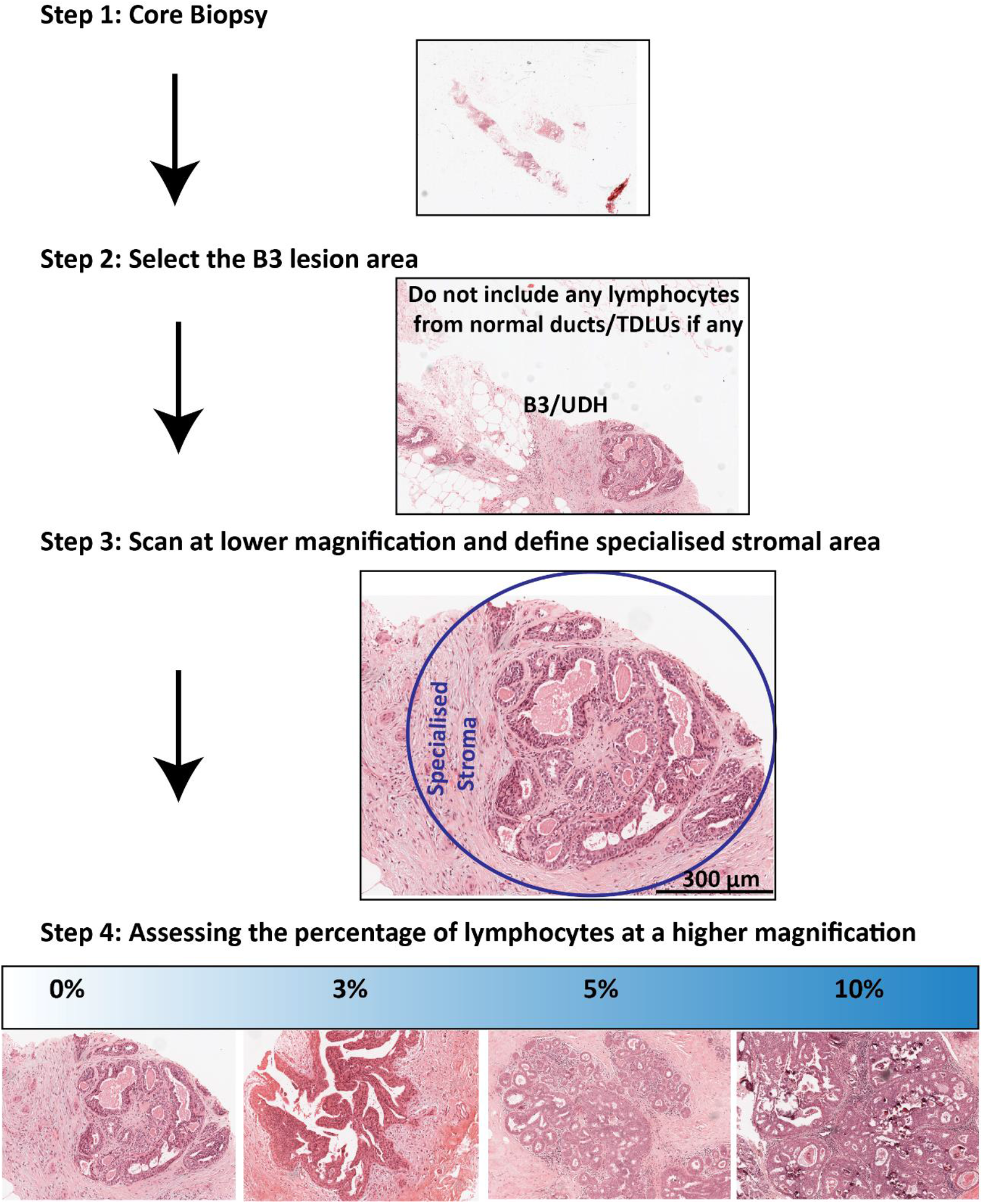
Step wise method for the evaluation of stromal lymphocytes of B3 lesions on H&E biopsy sections. This method of counting TILs from benign breast disease has recently been published by International Immuno-Oncological Working group (www.tilsinbreastcancer.org) (Rohan *et al* BCR 2021, 23:15).

Similarly, most cases of the validation cohort met the criteria for the lymphocyte assessment of B3 lesions (ductal lesions: n= 24 non-upgraded and 15 upgraded; papillary lesions: n= 43 non-upgraded and n=8 upgraded) (Supp Figure 2).

### Samples of upgraded ductal or papillary B3 lesions

Cases were considered upgraded (ductal or papillary) B3 when a diagnosis of B3 was made on core biopsy and DCIS/IBC was diagnosed on subsequent excision within a year after B3 diagnosis. Cases were considered non-upgraded when no DCIS/IBC was reported on subsequent excision within a year after B3 diagnosis.

### Samples of DCIS

Data for LG DCIS cases (n=19), intermediate grade (IG) DCIS (n=36) and high grade (HG) DCIS (n= 96) were derived from the previously published work [18], with new data from an additional six LG DCIS cases from the RMH, St Vincent’s, Peter MacCallum Cancer Centre [23].

### Ethics approval

This study was conducted under ethical approval from the Peter MacCallum Cancer Centre (HREC #12-64), (HREC#19/194), Melbourne Health (HREC# 2012.119), (HREC#2019.390), St Vincent’s Hospital (HREC #022/19), the North West-Greater Manchester Central Research Ethics Committee 15/NW/0685.

### Statistical analysis

GraphPad Prism v8 (San Diego, CA, USA) and R (v.4.0.2) were used to generate graphs and perform statistical tests as indicated. A p value of <0.05 was considered significant. An area under the curve (AUC) was created using a logistic regression model with lymphocytes count as a continuous variable adjusted by patient’s age as a non-linear term (9 cases were excluded due to missing data on patient’s age). C-stat and AUC were reported for this analysis. Since we have identified a threshold of stromal lymphocytes (≥5%) to predict upgraded cases, specificity and sensitivity were defined as follows: specificity was true negative tests divided by sum of true negative and false positive; sensitivity was true positive tests divided by sum of true positive and false negative.

## Results

We investigated the stromal lymphocytes in our discovery cohort of B3 lesions. This cohort consists of non-upgraded ADH (n=43), other non-ADH ductal B3 lesions (n=55) and papillary lesions (n=31). We compared these non-upgraded lesions to upgraded ductal (n=36) and papillary (n=9) lesions. The median age at diagnosis of non-upgraded patients was 55 (range 20-76) and upgraded patients was 58 (range 40-73) (p=0.14, Mann-Whitney t-test). Overall, most of these patients were diagnosed after routine mammogram imaging and were classified under category 2 or 3 regardless of the upgrade status at the time of diagnosis (Table 1). The retrospective cohorts in this study were diagnosed between 1995-2016 with a mammogram imaging category system (i.e. not BIRADS) whereby category 1 = no significant abnormality, category 2 = benign findings, category 3 = indeterminate/equivocal findings, category 4 = findings suspicious of malignancy and category 5 = malignant findings.

**Table 1.** Sample cohort.

| Characteristics | Non-upgraded B3 (ductal + papillary) | Upgraded B3 (ductal + papillary) | *P* value (test) |
| --- | --- | --- | --- |
| Median age at diagnosis (yr) (range) | 55 (20-76) (n=124) | 58 (40-73) (n=41) | 0.14 (MW) |
|  | *57 (42-78) (n=67)* | *61 (46-79) (n=23)* | *0.11 (MW)* |
| Imaging category 2 or 3 | 38/45 (84%) | 22/25 (88%) | 0.74 (FE) |
|  | *56/67 (84%)* | *17/23 (74%)* | *0.36 (FE)* |
| Imaging category 4 or 5 | 7/45 (16%) | 3/25 (12%) |  |
|  | *11/67 (16%)* | *6/23 (26%)* |  |

MW: Mann-Whitney test two-tailed, FE: Fisher Exact test two-tailed, cohort in *italic* represents validation cohort.

### Stromal lymphocytes were not frequently observed in non-upgraded other non-ADH ductal B3 lesions

The presence of stromal lymphocytes in the surrounding specialised stroma of 43 non-upgraded ADH ducts/foci of the experimental cohort was assessed. The presence of lymphocytes in non-upgraded ADH cases ranged from a score of 0-5%, with a median of 1% (Fig 2A, Supp File 1). The presence of stromal lymphocytes in other non-ADH ductal B3 lesions ranged from 0-5%, with a median of 0% (Fig 2A). When the presence of stromal lymphocytes was compared between non-upgraded ADH and non-upgraded other non-ADH ductal B3 lesions, there was a significant difference observed (p=0.0058, zero inflated negative binomial distribution) (Fig 2A). Sixteen non-upgraded ADH cases (16/43, 37.2%) had no lymphocytes (i.e. 0% lymphocytes) whereas 43 other non-ADH ductal B3 lesions were observed with no lymphocytes (43/55, 78.2%). Three ADH cases showed a maximum score of 5% (3/43, 7%). Ten other non-ADH ductal B3 cases had 1-2% (10/55, 18%) stromal lymphocytes present while two cases (2/55, 3.6%) had 5% stromal lymphocytes present.

**Figure 2.**
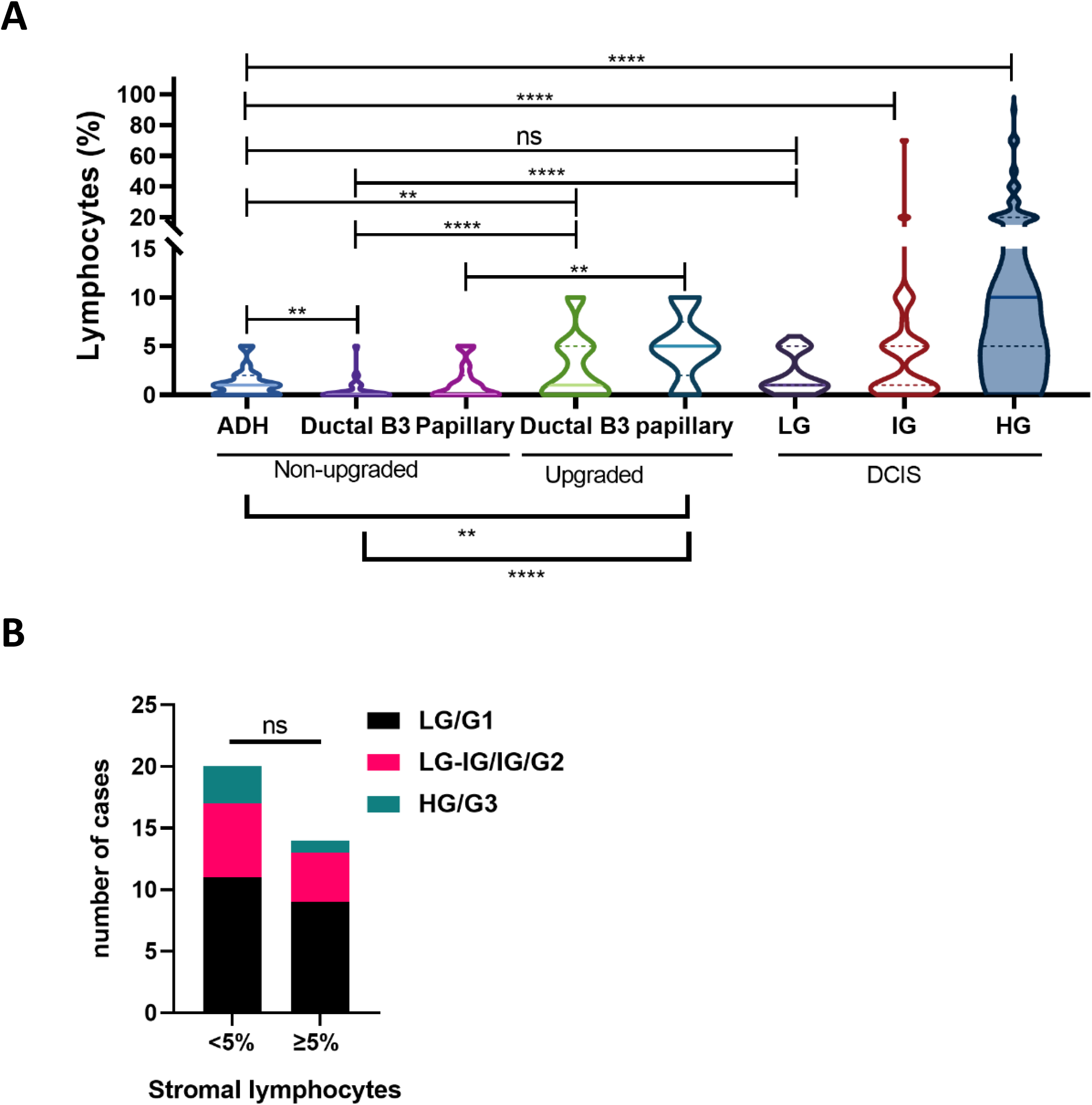
Predicting upgraded B3 from biopsies utilising total stromal lymphocytes (%) in discovery cohorts. A. The violin plot represents stromal lymphocyte counts (%) of non-upgraded ADH (n=43), other non-ADH ductal lesions (n=55), and papillary lesions (n=31) and compares to upgraded ductal (ADH and non-ADH B3, n=36) and papillary (n=9) B3 cases. The lymphocyte counts (%) of non-upgraded ADH was also compared to previously published low (n=25), inter (n=36) and high grade (n=96) DCIS (Hendry *et al* CCR 2017). *p<0.05; *** p<0.001; **** p<0.0001; zero-inflated negative binomial model. Solid line of violin plots represents median and dotted lines quartiles. B. Stromal lymphocytes (<5% or ≥5%) of upgraded B3 cases in the discovery cohort were compared to the grades of DCIS/IDC these B3 cases have been upgraded to. ns = not significant, Fisher Exact test two tailed.

We compared the presence of stromal lymphocytes of B3 and ADH cases with TILs of LG DCIS (n=25), IG DCIS (n=36) and HG DCIS (n=96) (most previously published) [18]. IG and HG DCIS showed a significant increase in total lymphocytes compared to that of ADH (p<0.0001, p<0.0001, respectively, zero inflated negative binomial model) (Fig 2A) with a median of 3% (range 0-70%) and 10% (range 0-90%), respectively, of stromal lymphocytes. Seven LG DCIS cases (7/25, 28%) had a maximum score of 5-6% while only five cases showed no lymphocytes.

### Upgraded B3 lesions showed significantly higher stromal lymphocytes than the non-upgraded in discovery cohorts

We then investigated the total stromal lymphocytes of the upgraded B3 cases, which includes all types of B3 (including ADH) in the discovery cohort (Supp File 1) with the aim of predicting upgrade from core biopsies. It is noteworthy that most upgraded cases were ADH, except one case of FEA and radial scar. The upgrade tumours spanned a range of grades and histological types (Supp File 1). Overall, 45% (20/45) of all upgraded B3 cases were upgraded to LG DCIS/G1 IBC (12/20 = pure LG DCIS), 22% (10/45) to LG-IG DCIS/IG DCIS/G2 IBC, 9% (4/45) to HG DCIS/G3 IBC and 7% (3/45) to lobular/papillary subtypes of cancer (Supp Figure 5). When the grades of DCIS/IBC are mixed, they are categorised under the highest grade of tumour observed. The grade of 17% (8/45) and estrogen receptor status of all DCIS/IBC cases were not available for the discovery cohort.

The presence of stromal lymphocytes in upgraded ductal B3 cases ranged from a score of 0-10%, with a median of 1% (Fig 2B, Supp File 1). Upgraded ductal B3 lesions had significantly higher stromal lymphocytes than the non-upgraded ADH and other ductal non-ADH B3 lesions (p=0.0058, p<0.0001, respectively, zero inflated negative binomial model) (Fig 2A). Four upgraded ADH cases showed the highest score of 10% stromal lymphocytes (4/36, 11%). Nine upgraded ADH cases had 5-9% stromal lymphocytes (9/36, 25%) while one-third of upgraded ductal B3 cases showed no stromal lymphocytes (0%).

We also investigated papillary lesions with the aim of predicting upgrades from core biopsies of discovery cohorts. Papillary lesions included in these discovery cohorts were both benign and atypical. It is noteworthy that unlike ADH/ductal B3, papillary lesions frequently did not meet the criteria to count lymphocytes (54% (46/86) of biopsies, Supp Figure 1). There was a significant difference between non-upgraded papillary lesions and upgraded (p<0.0001, zero inflated negative binomial model) with regards to total lymphocytes with a median of 0% (range 0-5%) *vs* 5% (range 0-10%), respectively (Fig 2A).

Degnim *et al* [28] showed that patient age was one of the risk factors for subsequent cancer after a diagnosis of atypical hyperplasia. Therefore, for application in a clinical setting, we developed a logistic regression model with patient age and lymphocyte count as variables, with an AUC of 0.77 (Supp Figure 6). If we categorise lymphocyte count as high (≥5%) or low (<5%) adjusted to patient age and type of B3 lesion (ductal or papillary), the AUC remained similar (0.72).

There was no significant correlation between the presence of ≥5% stromal lymphocytes on upgraded biopsies and the grade of DCIS/IBC diagnosed on surgical excision (p=0.89, Fisher exact test two-tailed). For example, of the 20 biopsies upgraded to LG DCIS/G1 IBC 11 had <5% lymphocytes and 9 had ≥5% lymphocytes. Similarly, for higher grade carcinomas 9/14 biopsies showed <5% lymphocytes and 5/14 showed ≥5% (Fig 2B).

### Validation of stromal lymphocytes as a clinical tool for diagnosis and predicting upgraded B3 lesions of all types

Since our discovery cohort showed a significant difference between upgraded and non-upgraded B3 cases, we then aimed to validate this as an immune biomarker in another patient cohort using the ≥5% lymphocytes threshold determined in the discovery cohort. The total number of cases in this validation cohort was 67 non-upgraded B3 and 23 upgraded, which includes ADH, non-ADH ductal other B3 as well as papillary lesions. These were ascertained as a consecutive cohort of all B3 diagnoses in a population BreastScreen program (NWBS) over the period of 2004-2013.

The median age at diagnosis of patients from the validation cohort was 57 (range 42-78) for non-upgraded and 61 (range 46-79) for upgraded (p=0.11, Mann-Whitney t-test). Most cases in the validation cohort were classified under mammogram imaging category 2 or 3 (benign/indeterminate) regardless of the status of upgrades (Table 1). The grades of the upgraded carcinomas were unknown for most cases (14/23: 61%). In the validation cohort the ductal B3 non-upgraded lesions comprised ADH as well as other low risk B3 such as CCL and FEA. Only eleven out of 24 non-upgraded cases (46%) showed no lymphocytes as opposed to the discovery cohort in which 78% other non-ADH ductal B3 cases showed no lymphocytes.

There was a significant difference observed with regards to the stromal lymphocytes of ductal and papillary B3 cases in the validation cohort between upgraded (median 0%) versus non-upgraded (median 5%, p<0.0008, p=0.023, zero-inflated negative binomial model, Fig 3A, 3B). Of cases with <5% lymphocytes, 16% were upgraded, compared to 57% of cases with ≥5% lymphocytes.

**Figure 3.**
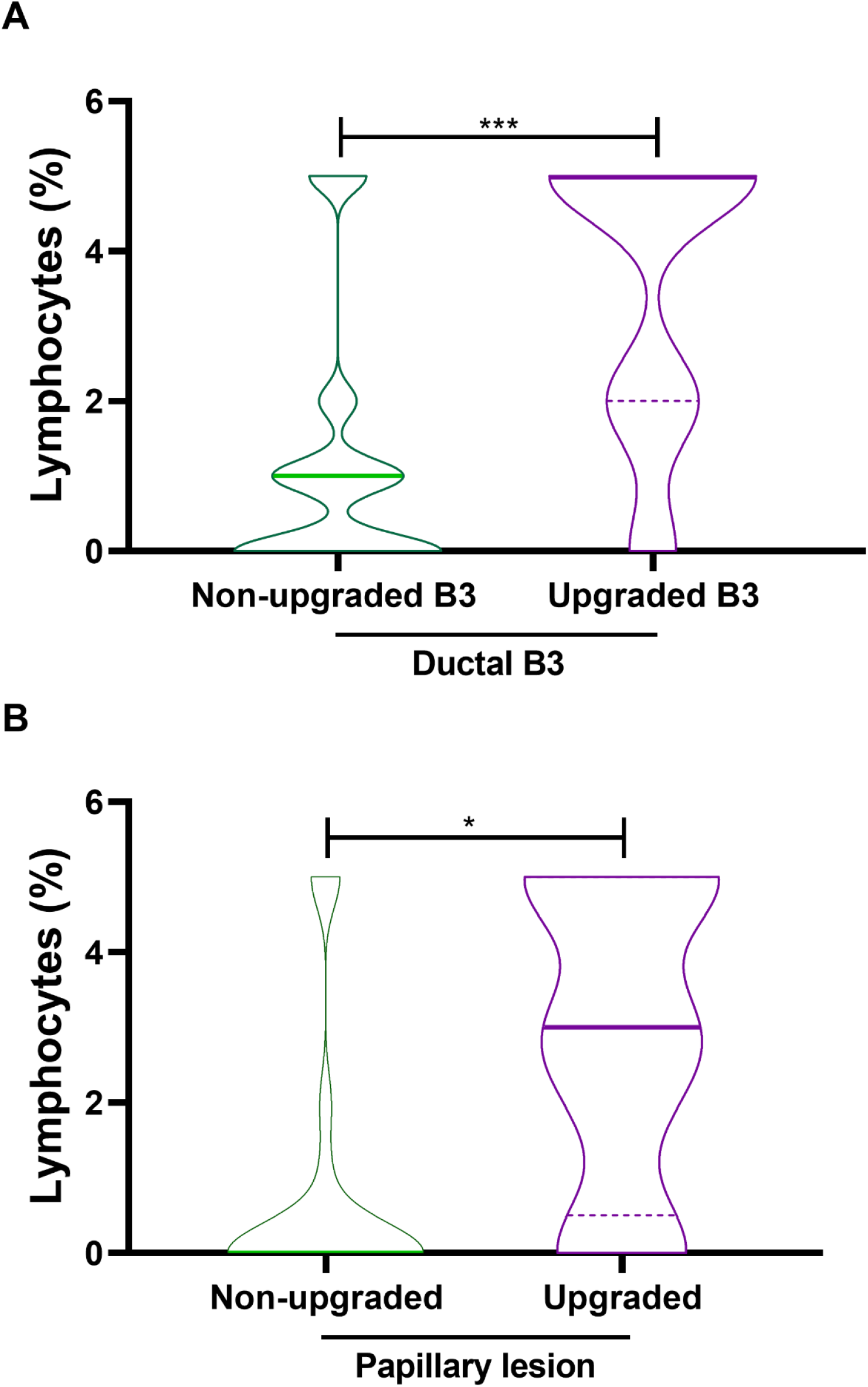
Predicting upgraded B3 from biopsies utilising total stromal lymphocytes (%) in validation cohort. A. The violin plot represents stromal lymphocytes counts (%) of non-upgraded ductal B3 includes ADH and non-ADH ductal lesions (n=23) and compares to upgraded ductal (n=15) B3 cases. This dataset excludes one outlier non-upgraded ductal B3 case (NWBS39) which shows 15% total lymphocytes on biopsy but only ALH was found on subsequent surgical excision (Supp File1). B. The violin plot represents stromal lymphocytes counts (%) of non-upgraded papillary (n=43) and compares to upgraded papillary lesions (n=8). *p<0.05; *** p<0.001; zero-inflated negative binomial model. Solid line of violin plots represents median and dotted lines quartiles.

### Potential biomarker to predict upgraded B3 lesions from core biopsies utilising ≥5% stromal lymphocytes

Taken all together, the sensitivity and specificity of predicting upgraded B3 cases from core biopsies utilising ≥5% stromal lymphocytes are 42% and 93%, respectively in our discovery cohorts. Similarly, the sensitivity and specificity in our validation cohort was 52% and 87%, respectively. This data suggests that the presence of ≥5% stromal lymphocytes on B3 biopsies indicates increased risk of upgrade. The logistic regression model developed with the discovery cohort was then applied on the validation cohort with the same parameters. The C-index or AUC of the validation cohort was 0.81, similar to the discovery cohort and if we categorise lymphocyte count as high or low and adjust for type of lesion and patient age, the AUC of this cohort was 0.73.

## Discussion

Many efforts to identify predictive biomarkers of upgraded B3 lesions have failed because of either a lack of specificity: the “low risk” group still has a considerable risk of upgrade [29], the predictive feature is not reproducible [30, 31] or the feature is only prognostic after full excision [28, 32]. Overall, predictive features are inconsistent across studies and many of the features require highly experienced pathologists and radiologists to interpret the available data.

Cases that are histologically B3 on biopsy but with a mammographic imaging category indicating malignant findings are not problematical clinically due to the clear clinical guidelines for these cases to be surgically excised. On the other hand, B3 cases on biopsy with mammographic findings equivocal/indeterminate or suspicious for malignancy cannot be ruled out for co-existing cancer only based on imaging, leading to potentially unnecessary surgeries. Most of our B3 biopsies are recorded as either indeterminate (category 3) or suspicious (category 4) on mammogram imaging. Here we developed a predictive marker for all types of B3 lesions based on total stromal lymphocytes as well as patient age and reported for the first time a predictive biomarker of upgrades that would be reproducible and cost effective anywhere in the world. Here we have shown that the presence of ≥5% lymphocytes on biopsy indicates a lesion that is more likely to be upgraded with a high specificity of >85% and an AUC of >0.75. These patients should not be considered for omission of surgical removal of the lesion. In contrast, absence of lymphocytes indicates the B3 lesion could be more likely to be benign. These patients may consider surveillance over immediate surgery, reducing unnecessary surgeries. It was shown that for example, CCL and FEA if correctly diagnosed carry a low risk of upgrade [7–9, 33].

It is important to keep in mind that the distinction between ADH and LG DCIS is often only based on the size of the lesion. Thus, LG DCIS may not be diagnosed at biopsy due to insufficient sampling of the lesion leading to a preliminary diagnosis of ADH. Therefore, it is debatable whether these cases should be considered “upgrades”. Nonetheless, the clinical consequence of a diagnosis of LG DCIS is different from that of ADH, although this may change should surveillance and not surgery become acceptable for LG DCIS. In this study, we included all cases even when upgraded to LG DCIS, because any predictive biomarker to enable safe observation will be desirable in clinical settings at this stage.

The logistic regression model using lymphocyte count and patient age as variables created an AUC of 0.729 and 0.732 in our discovery and validation cohorts, respectively. A risk score was recently developed by Lustig *et al*. [34] to predict upgraded ADH after using a logistic regression model to select variables. However, the score was based on 5 variables, such as size of multi-modal imaging and the presence of “DCIS-like” or > 1 high risk lesions on core biopsies. Our study on the other hand showed a high specificity of predicting any B3 lesions using simpler variables with a high reproducibility.

However, using lymphocytes alone as a marker would under-diagnose approximately 50% of patients with upgrade. Nonetheless it was notable that 10/23 upgrades in the ductal B3 discovery cohort with <5% lymphocytes were small LG or IG DCIS, which potentially are lesions with low risk of progression to more serious pathology if left untreated until the next routine screen. A further limitation is that the ability to apply this method to papilloma biopsies was hampered, with not quite half of the biopsies carrying sufficient stroma to score. There was no significant difference between the drop-out rate for upgrades (53%) and non-upgrades (59%)

A few key individual immune cell types were previously investigated in a large cohort of benign breast disease including ADH [35], however, this was never explored for upgrades. The biology of a higher number of lymphocytes and individual immune cells of upgraded cases are yet to be explored, but could reflect the altered microenvironment caused by the presence of nearby carcinoma.

The strength of our study is extensive pathological review by experienced pathologists to reconfirm the diagnosis of types of B3 lesions on biopsies with four independent patient cohorts. To our knowledge, this is the largest cohort to date analysed for lymphocytes of B3 lesions, which identified lymphocytes alone as a predictive biomarker of upgrades with a high specificity. This method of counting lymphocytes (or TILs for IBC and DCIS) is already accepted as part of the routine diagnostic procedure at many hospitals worldwide. The implementation of TILs has been informative for prognosis of IBC patients. Here we showed evidence of stromal lymphocytes of B3 biopsies combining patient age being predictive of upgraded cases, which are a clinical problem that must be addressed before de-escalating treatment for B3 lesions.

## Acknowledgements

We thank A/P Prue Hill (St Vincent’s Hospital, Melbourne) (retired) and Maria Bisignano (Melbourne Health Pathology) for the review of StVs B3 cases and for coordinating NWBS cases, respectively. We thank the Nottingham Health Science Biobank and Breast Cancer Now Tissue Bank for the provision of tissue samples.

## Authors’ Contributions

Conception and design: TK, KLG; providing access to clinical samples: GBM, HF, SJ, EAR, SBF, EP; TIL counts: TK, SH, JMP; Performing experiments, ethics approval, pathological reviews, acquisition of data, analysis and interpretation of data: TK, SH, JMP, LT, KLG, SJ, MT, EP, NR, AM, GBM, IGC; Statistical support: MWJ; Identification of cases: SH, HK, KE, KLG, DB, EH, SJ, IMM, DJB, EP; Drafted the manuscript: TK, KLG. All authors read and approved the final manuscript; overall study supervision: KLG and IGC.

---

**Supp Figure 1.**
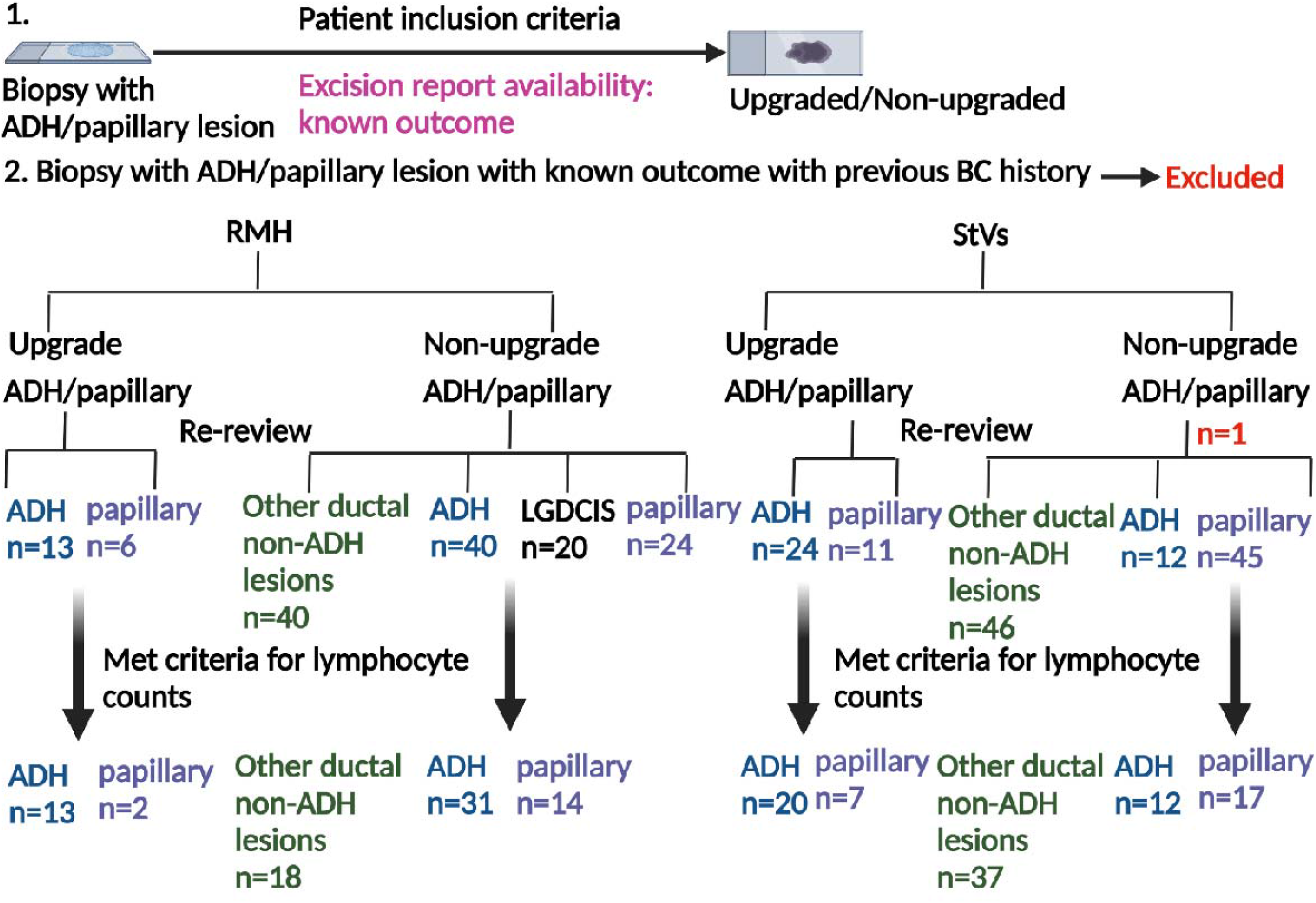
Patient inclusion and selection criteria of discovery cohorts. RMH: Royal Melbourne Hospital; StVs: St Vincent’s Hospital; BC: Breast cancer; ADH: atypical ductal hyperplasia; LGDCIS: Low grade ductal carcinoma in situ. There were additional 3 upgraded ADH cases from Nottingham cohort, UK included in this study (not shown here).

**Supp Figure 2.**
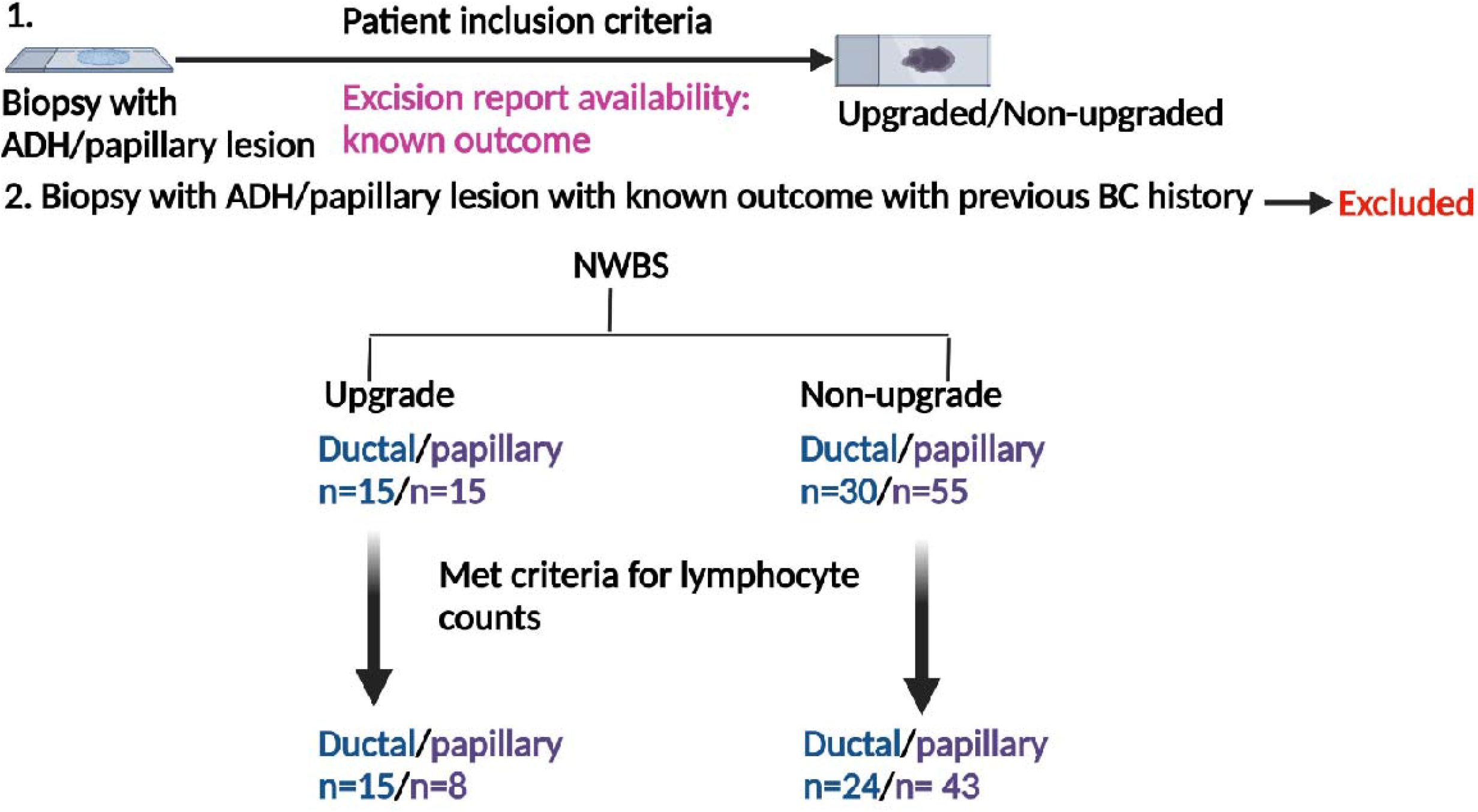
Patient inclusion and selection criteria of validation cohort. NWBS: North-west Breast Screen; BC: Breast cancer.

**Supp Figure 3.**
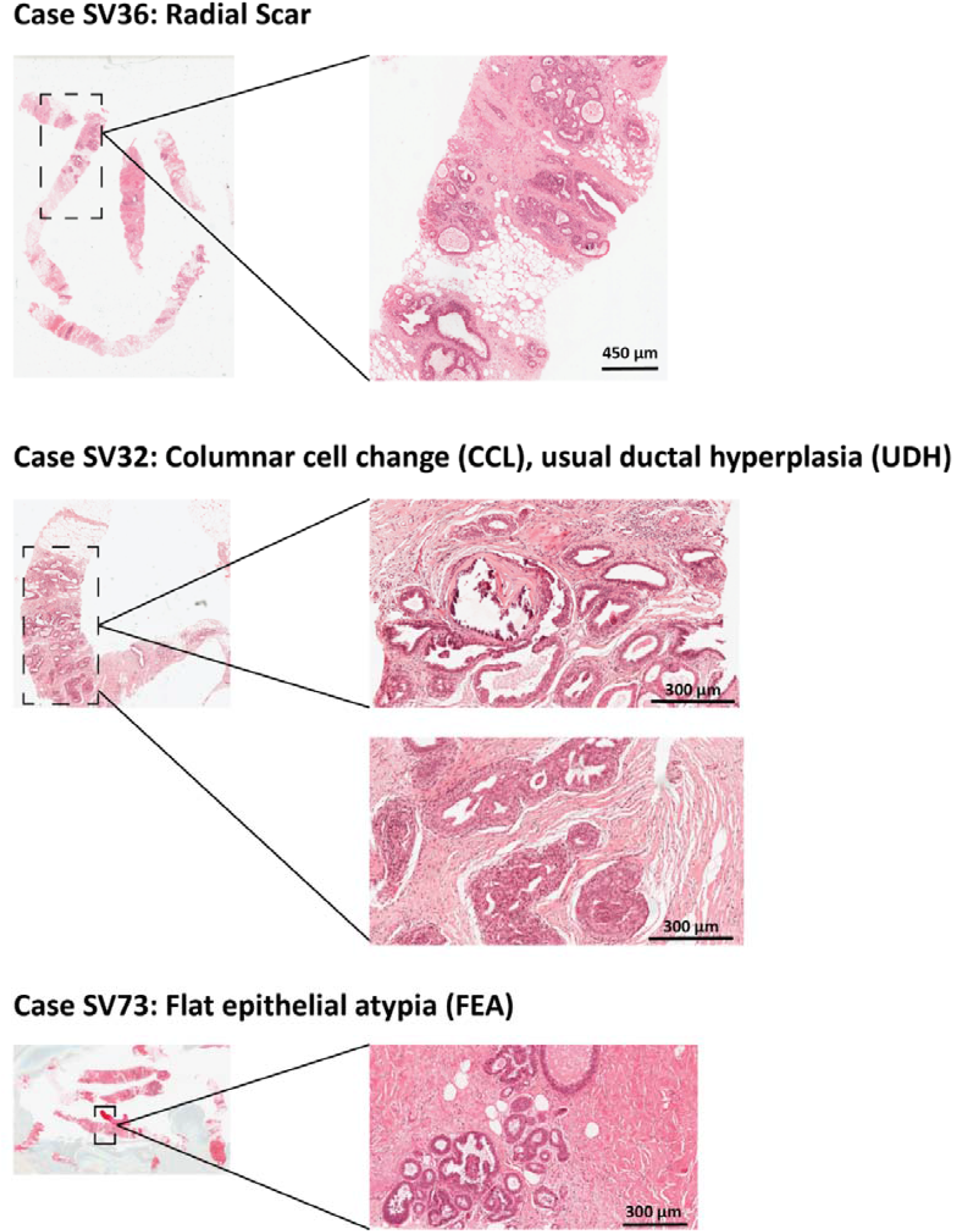
Examples of other ductal lesions that were not confirmed as ADH. These non-ADH ductal B3 lesions include radial scar, columnar cell change (CCL), flat epithelial atypia (FEA) or usual ductal hyperplasia (UDH).

**Supp Figure 4.**
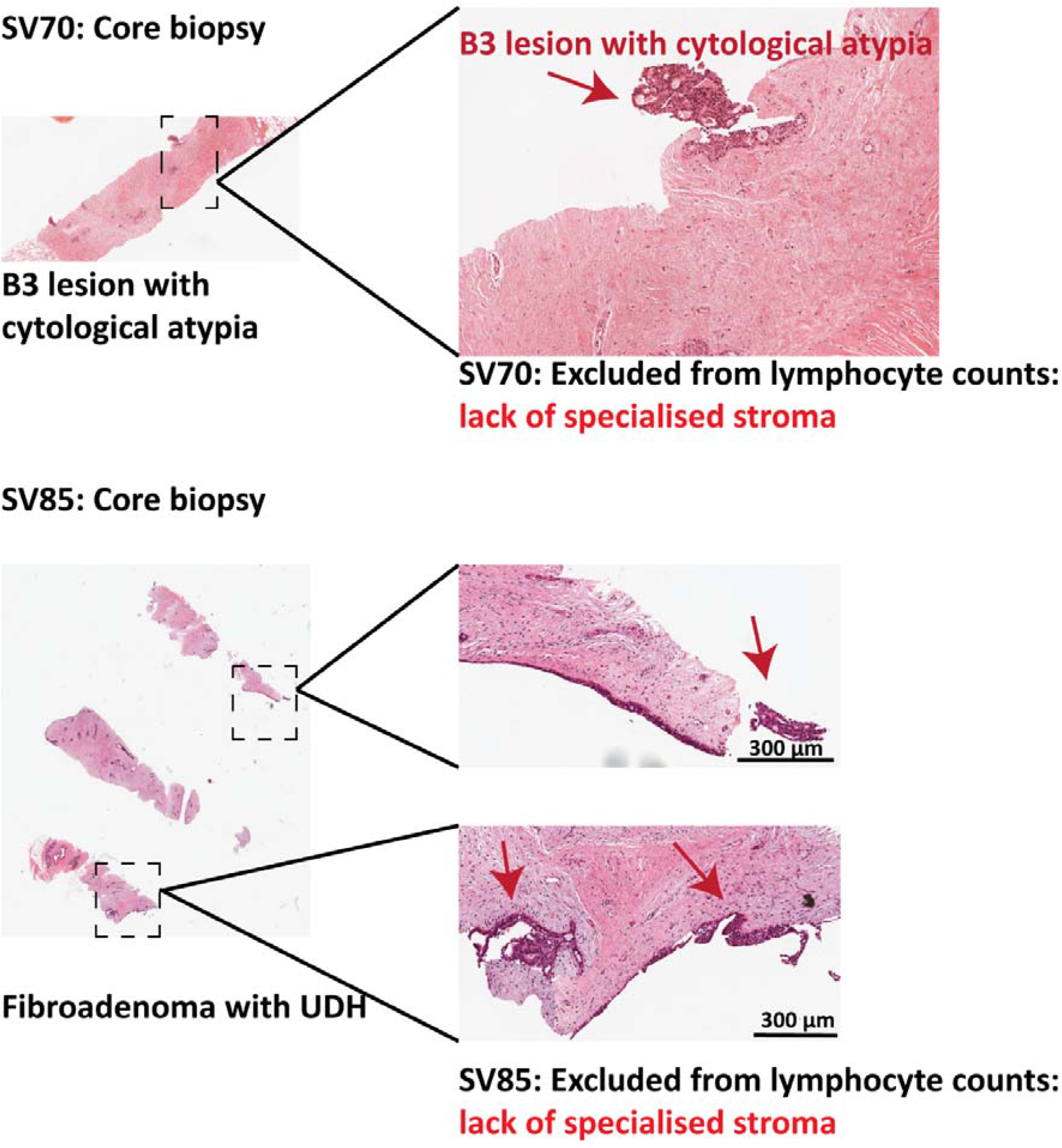
Examples of excluded cases from analysis due to lack of specialised stroma on B3 biopsies.

**Supp Figure 5.**
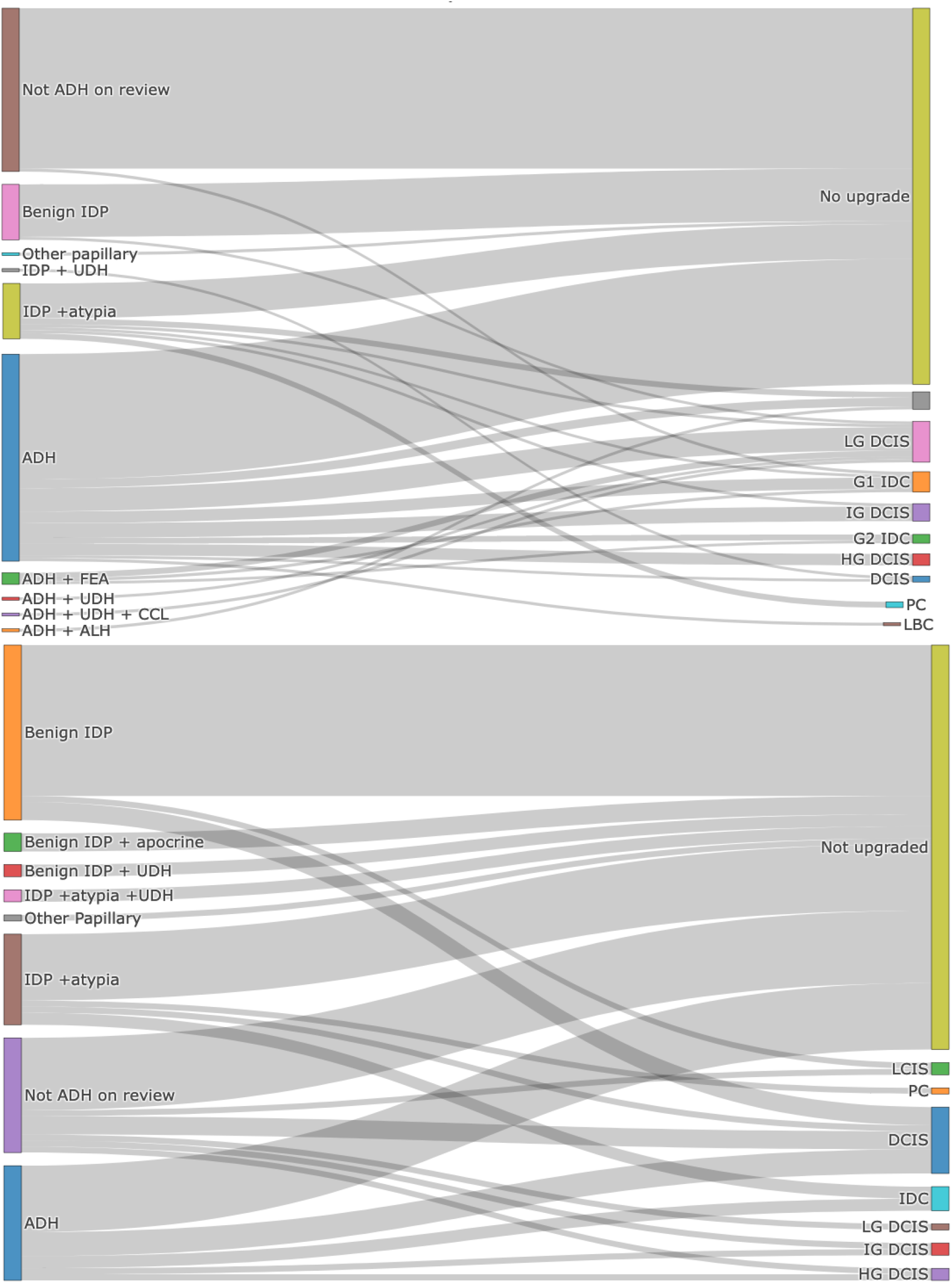
Sankey diagrams. The discovery cohort (top) and validation cohort (bottom) of upgraded and non-upgraded ADH, non-ADH ductal lesions, papillary lesions as well as grades and subtypes of DCIS/cancer they have been upgraded to (when available).

**Supp Figure 6.**
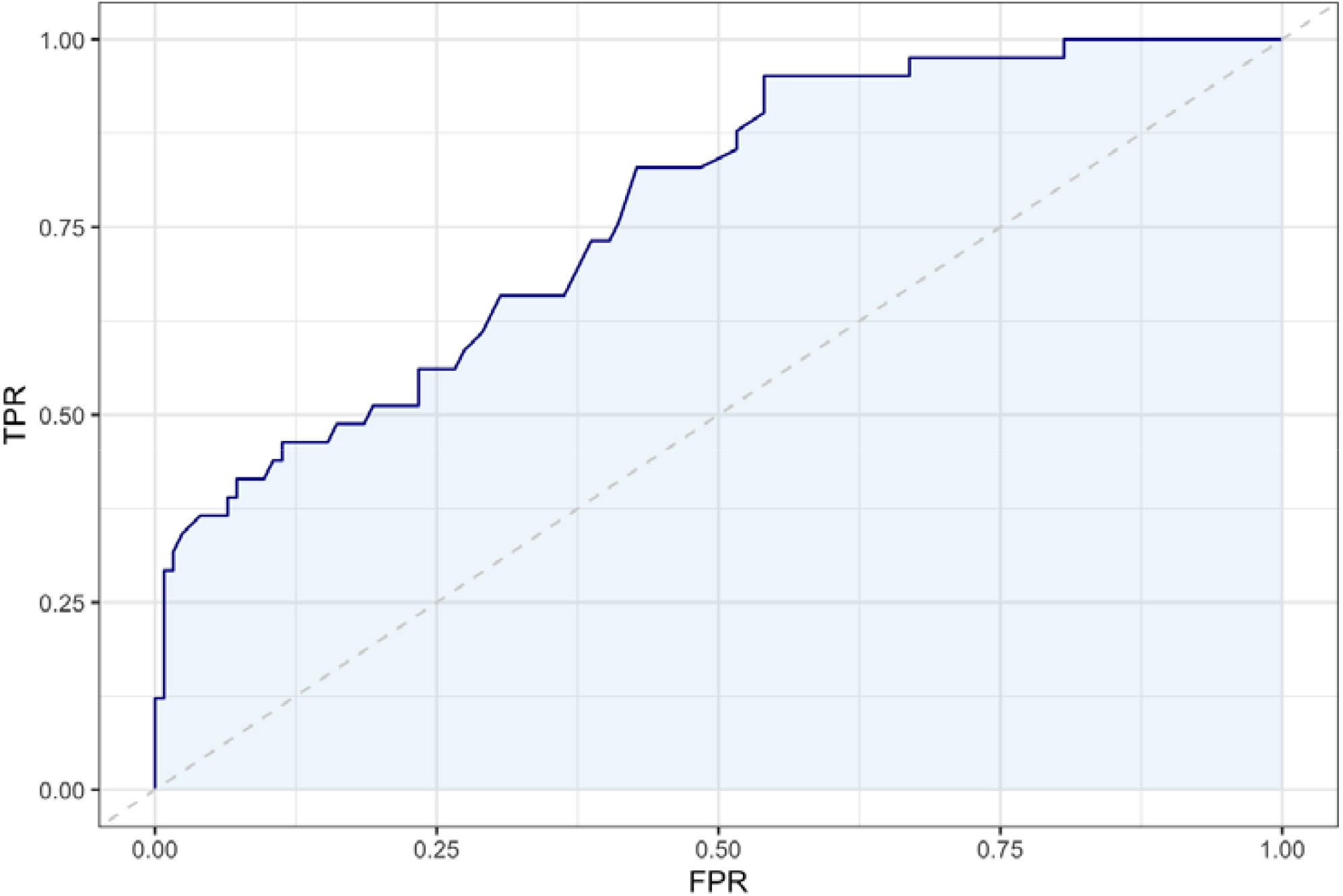
Area under the curve for discovery cohort. In our discovery cohort using lymphocyte count (as a continuous variable) and age as variables (n=165/174, n=9 excluded due to missing data on patient’s age), logistic regression model was developed and an AUC was created. Here AUC=0.77. X axis: TPR= True positive rate and Y axis: FPR= False positive rate. This curve was generated using logistic regression model with lymphocytes count adjusted to patient’s age as a non-linear term.

